# Experimentally-induced encoding variability influences mnemonic discrimination: Evidence from human behavioral data and global matching models

**DOI:** 10.64898/2026.02.10.705080

**Authors:** Derek J. Huffman, Leslie Rollins, Morgan Carter, Chayse A. Cotton, Katelyn B. Cockrell, Elena Rezac, Mai Khanh Tran

## Abstract

Computational models and neurobehavioral data suggest that encoding variability affects forced-choice mnemonic discrimination. Here, we experimentally manipulated encoding variability on the forced-choice Mnemonic Similarity Task by varying stimulus repetitions during encoding. We first generated predictions from a global matching model. Behavioral data supported all predictions. Across most conditions, repetitions consistently enhanced mnemonic discrimination; however, when encoding variability was induced by 3-repetitions of the original version of the non-corresponding lure and 1-repetition of the target during learning, individuals exhibited increased interference. These findings provide further insight into theories of human memory, especially the effect of stimulus repetition on mnemonic discrimination.

---

Prominent theories argue that pattern separation is a core mnemonic function of the hippocampus (e.g., Norman & O’Reilly, 2003; Treves & Rolls, 1992; Yassa & Stark, 2011). The Mnemonic Similarity Task [MST] was developed to test how well participants can disambiguate similar experiences in memory (Stark et al., 2013, 2015, 2019). In forced-choice versions of the MST (e.g., Huffman & Guan, 2025; Huffman & Stark, 2017), targets (A) are paired with unrelated foils (A-X), corresponding lures (i.e., a lure similar to the target; A-A’), or non-corresponding lures (i.e., a lure similar to a different item; A-B’). Performance is generally best on A-X, intermediate on A-A’, and worst, though better than chance, on A-B’ (Huffman & Guan, 2025; Huffman & Stark, 2017; Rollins et al., 2019, 2024; cf. Migo et al., 2009).

Evidence from computational modeling, behavioral data, and psychophysiological data suggests that encoding variability can account for the A-A’ *>* A-B’ test format effect. A global matching model, MINERVA 2, assumes that memories are composed of distributed representations of stimulus features (Hintzman, 1984, 1988). Encoding variability is defined by variability in how many features are encoded for each stimulus presentation. MINERVA 2 and other global matching models show that increased encoding of the original B item (lure) relative to the A item (target) can lead to increased interference on the A-B’ test format (Hintzman, 1988; Huffman & Guan, 2025; Huffman & Stark, 2017). Supporting the encoding variability predictions, empirical studies have shown that errors on the A-B’ test format were associated with more visual fixations (Rollins et al., 2019; 2023) and a higher amplitude ERP response (Rollins et al., 2024) during learning to the original version of the lure (B) relative to the target (A).

We argue that stimulus repetitions can be employed as a means of experimentally manipulating encoding variability. When viewed through this lens, we can reinterpret past findings and make several interesting new predictions via computational modeling. There is consensus that repetition improves the discrimination of targets and unrelated foils, but there is debate about whether repetition enhances (Loiotile & Courtney, 2015; Zhang & Hupbach, 2020) or impairs (Reagh & Yassa, 2014) mnemonic discrimination between targets and similar lures. Reagh and Yassa (2014) found that the lure discrimination index was paradoxically worse with 3 repetitions vs. 1 repetition. They argued that their results support Competitive Trace Theory (CTT; Yassa & Reagh, 2013) in which separate, similar memory traces are stored for each repetition, which they argue would result in semanticization. However, subsequent research using signal detection theory (SDT) found that 3 repetitions improved lure discrimination (Loiotile & Courtney, 2015; Zhang & Hupbach, 2020). We aimed to further test the predictions of CTT (Reagh & Yassa, 2014) vs. global matching models (Huffman & Guan, 2025) by combining novel test formats in the forced-choice MST with computational modeling.

Sixty-six young adults (*M* = 20.0 years, 44 females, 22 males) completed a modified version of the forced-choice MST (Supplemental Material). We selected 350 pairs of colored pictures from the Stark Lab’s stimulus set (https://faculty.sites.uci.edu/starklab/mnemonic-similarity-task-mst/). To counterbalance which stimuli were used as targets, lures, and foils across participants, we separated stimuli into 14 sets of 25 pictures with five pictures from each mnemonic similarity bin established in previous research (e.g., Stark et al., 2013, 2015, 2019). E-Prime software (Psychological Software tools, Inc., Pittsburgh PA) presented stimuli in a random order and collected behavioral responses. Participants incidentally encoded 600 items (i.e., 150 were presented once, 150 were presented thrice) by making an indoor/outdoor judgment. Each picture was shown for 2000 ms and preceded by a 500 ms fixation cross. At retrieval, participants completed 200 two-alternative forced-choice mnemonic discrimination trials across eight conditions (Figure 1). Each condition included 25 trials. Targets (A) and the original version of lures (B) were either encoded with 1-repetition (A1x-X, A1x-A’, A1x-B’1x) or 3-repetitions (A3x-X, A3x-A’, A3x-B’3x). We experimentally manipulated encoding variability by assessing performance on A-B’ trials for which there was an unequal number of repetitions for the A and B items during learning (i.e., A3x-B’1x, A1x-B’3x).

**Figure 1.**
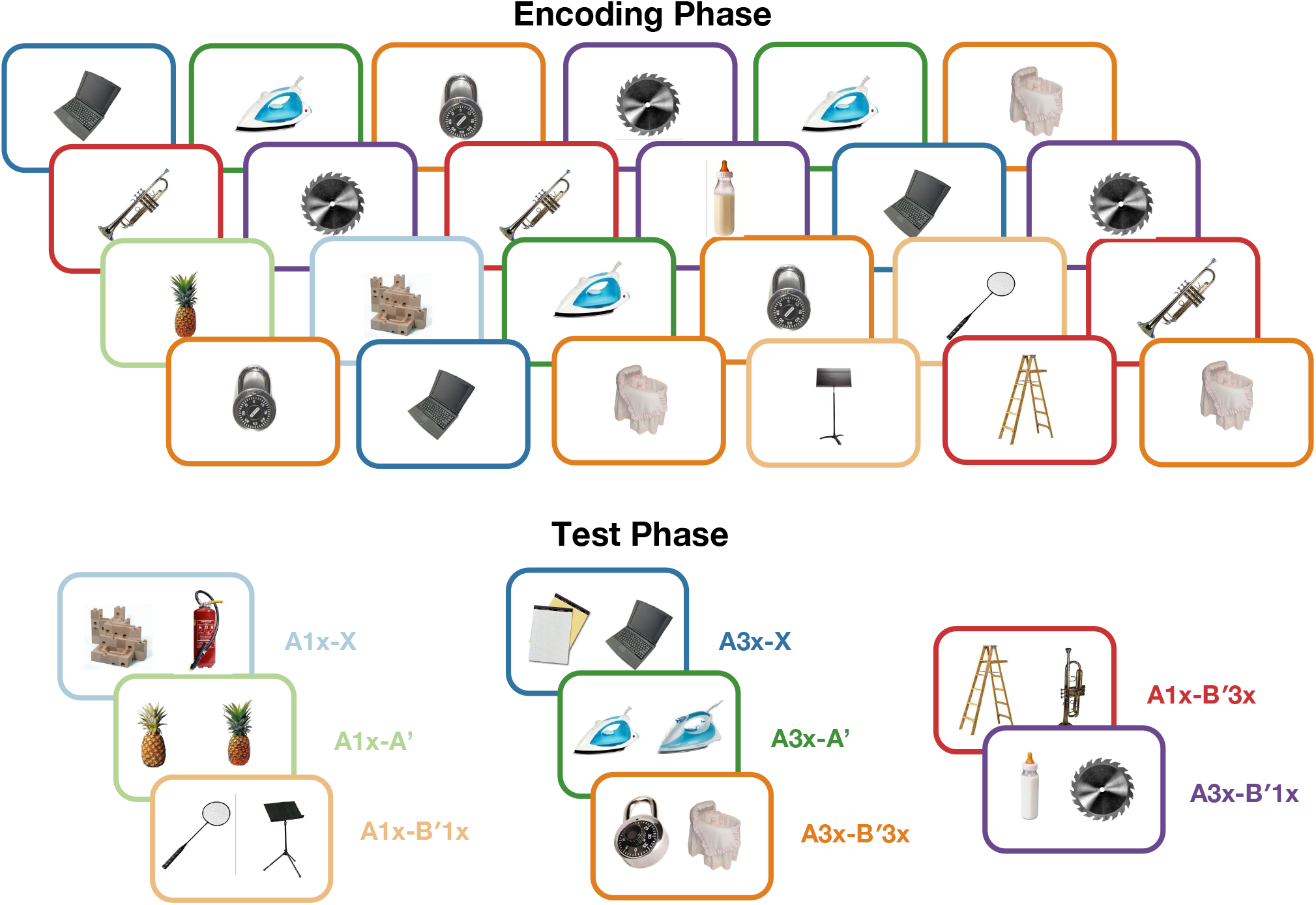
: Depiction of the behavioral experiment. Participants completed a modified version of the forced-choice MST. During the test phase, targets (A) were either paired with unrelated foils (A-X), corresponding lures (i.e., a lure similar to the target; A-A’), or non-corresponding lures (i.e., a lure similar to a different item; A-B’). Targets (A) and the original version of lures (B) were either encoded with 1-repetition (A1x-X, A1x-A’, A1x-B’1x) or 3-repetitions (A3x-X, A3x-A’, A3x-B’3x). We experimentally manipulated encoding variability by assessing performance on A-B’ trials for which there was an unequal number of repetitions for the A and B items during learning (i.e., A3x-B’1x, A1x-B’3x).

Prior to conducting our behavioral experiment, we generated predictions from a global matching model, MINERVA 2, about performance as a function of repetitions and test format (Figure 2A and Supplemental Material). MINERVA 2 predicted that repetitions would improve memory performance on the A-X, A-A’, and A-B’ test formats (see predictions 1-3 and Figure 2A below). Likewise, MINERVA 2 predicted that there would be a test format effect such that A-X *>* A-A’ *>* A-B’, which is also consistent with previous research (see Figure 2A and Huffman & Guan, 2025; Huffman & Stark, 2017). A key novel aspect to our computational modeling work and behavioral experiment was our experimentally-manipulated encoding variability conditions (i.e., A3x-B’1x, A1x-B’3x). We operationally defined the 3-repetition A-B’ condition (A3x-B’3x) as having greater chances for encoding features vs. the 1-repetition A-B’ condition (A1x-B’1x). Thus, comparisons of trials with unequal repetitions (i.e., A3x-B’1x, A1x-B’3x) allowed us to test the effect of encoding variability on mnemonic discrimination. MINERVA 2 generated an interesting set of predictions (Figure 2A), and, importantly, all of these predictions were supported by our behavioral results. Below we provide each of the novel MINERVA 2 predictions followed by the results of the corresponding behavioral analysis (note: all of the *β*s are positive, indicating that the results follow the expected pattern, and all *ps <* .05). In, summary, MINERVA 2 predicted better performance when targets (A) were presented with 3-repetitions than 1-repetition (Predictions 1-6) and better performance when the original version of the non-corresponding lure (B) was presented with 1-repetition than 3-repetitions (Predictions 6-8; note: the labels indicate whether the CTT prediction is: ^1^=the same, ^2^=opposite, ^3^=unclear, Supplemental Material):

**Figure 2.**
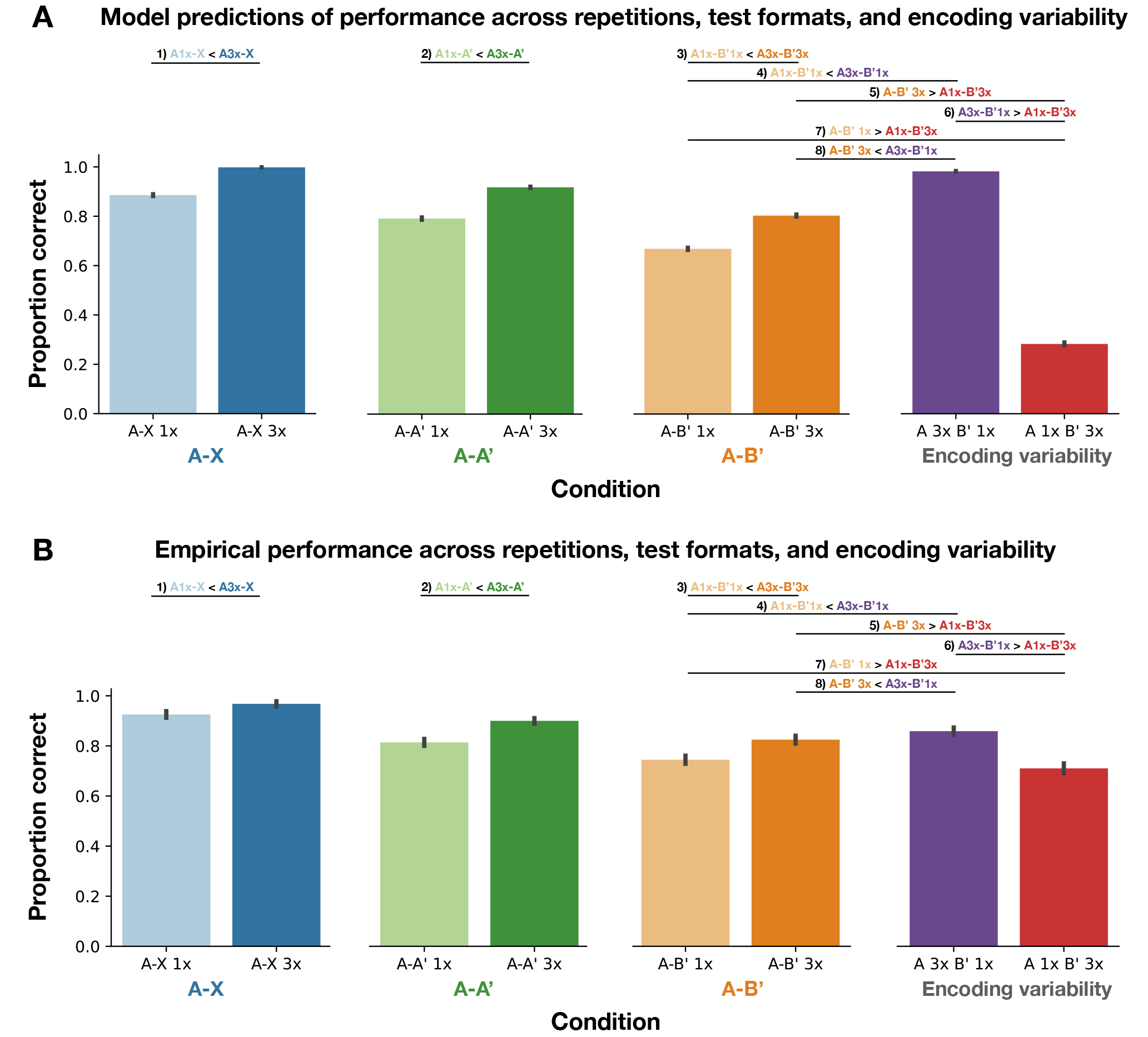
: The empirical data support MINERVA 2’s predictions of differences in performance across the test formats. Top: MINERVA 2 predicts better performance when targets (A) are presented with 3-repetitions than 1-repetition (Predictions 1-6) and when the original version of the non-corresponding lure (B) was presented with 1-repetition than 3-repetitions (Predictions 6-8). MINERVA 2 also predicted a test-format effect: A-X > A-A’ > A-B’ (as can be seen visually). Bottom: The empirical data supported all of the MINERVA 2 predictions of performance as a function of test format, repetitions, and encoding variability (all *ps* < .05; see main text). The data for the plots indicate the raw means and 95% standard errors but we used a binomial model for analysis.

1. Better performance^1^ on A3x-X vs. A1x-X, *β* = 1.04, *χ*^2^(1) = 35.15, *p <* .001.
2. Better performance^2^ on A3x-A’ vs. A1x-A’, *β* = 0.76, *χ*^2^(1) = 53.12, *p <* .001.
3. Better performance^2^ on A3x-B’3x vs. A1x-B’1x, *β* = 0.51, *χ*^2^(1) = 33.40, *p <* .001.
4. Better performance^3^ for A3x-B’1x vs. A1x-B’1x, *β* = 0.77, *χ*^2^(1) = 71.90, *p <* .001.
5. Better performance^3^ for A3x-B’3X vs. A1x-B’3x, *β* = 0.71, *χ*^2^(1) = 66.43, *p <* .001.
6. Better performance^3^ for A3x-B’1x vs. A1x-B’3x, *β* = 0.97, *χ*^2^(1) = 116.10, *p <* .001.
7. Better performance^1^ for A1x-B’1x vs. A1xB’3x, *β* = 0.19, *χ*^2^(1) = 5.29, *p* = .021.
8. Better performance^1^ for A3x-B’1x vs. A3x-B’3x, *β* = 0.27, *χ*^2^(1) = 7.68, *p* = .006.

More specifically, we determined whether the results from the behavioral experiment supported the predictions of MINERVA 2 on the traditional forced-choice test formats: A-X, A-A’, and A-B’ with 1- and 3-repetitions within each test format. We set up a model with the following contrasts: 1) a repetition contrast (+0.5 for the 3 repetitions and -0.5 for the 1 repetitions), 2) a test-format contrast (+1 for A-X, 0 for A-A’, and -1 for A-B’), 3) a repetition-contrast × test-format-contrast interaction. Here, we used binomial models within a mixed-model framework as follows using the mixed function from the afex package in R and a binomial model (our random effects structure included random intercepts as a function of participants and random slopes for the test-format contrast):

~~~
model <-mixed(
   ProportionCorrect ∼ Repetitions * TestFormat + (TestFormat | ParticipantID),
   data=data, method=‘LRT’, family=binomial, weight=data$n)
~~~

where `ProportionCorrect` is the proportion correct for a given format, `Repetitions` is a contrast of the predicted 3-repetitions > 1-repetition effect (see above), `TestFormat` is a contrast of the predicted A-X > A-A’ > A-B’ test format effect (see above), `weight=data`$`n` is a vector of the number of trials within each condition (25), Consistent with the predictions of MINERVA 2, we observed a significant effect of the repetitions contrast (*β* = 0.74, *χ*^2^(1) = 113.70, *p <* .001) and the test-format contrast (*β* = 0.91, *χ*^2^(1) = 102.15, *p <* .001). Moreover, we observed a significant repetitions-contrast × test-format-contrast interaction (*β* = 0.20, *χ*^2^(1) = 5.67, *p* = .017). Overall, these findings match the predictions of MINERVA 2 and suggest that repetitions enhance memory, there is a A-X *>* A-A’ *>* A-B’ test-format effect, and the benefit of repetitions may differ as a function of test format. To probe the interaction, we conducted a follow-up pairwise analysis of estimated marginal means (using emmeans within R; here, we used a categorical version of the Repetitions and Test Format variables for easier interpretation of the effects); this analysis revealed that the effect size for repetition was largest for the A-X format (odds ratio = 2.49, *p* < .0001) followed by the A-A’ format (odds ratio = 2.15, *p* < .0001) and smallest for the A-B’ format (odds ratio = 1.68, *p* < .0001). These findings are further supported by the positive *β* for the interaction term, thus suggesting a supra-additive effect of repetitions across the test format contrast. While this finding is admittedly non-intuitive by visually inspecting Figure 2B, it is driven by the non-linear nature of the binomial model, which causes correspondingly larger changes in magnitude as it approaches the ceiling of perfect performance (i.e., the logit scale; Supplemental Figure 1). We note that we performed a follow-up analysis on the predictions from MINERVA 2 and found that it also predicted an interaction (see Supplemental Material).

We further probed the nature of the repetition and test-format effects in follow-up tests within each condition. To test whether the effect of repetition was consistent across all of the test formats (see Predictions 1-3 from MINERVA 2), we defined our models as follows:

~~~
model <-mixed(ProportionCorrect ∼ Repetitions + (1 | ParticipantID),
   data=data, method=‘LRT’, family=binomial, weight=data$n)
~~~

where the variables are defined as above and the model includes random intercepts for participants (more complex random effects models failed to converge). We found a significant effect of the repetitions contrast for A-X (*β* = 1.04, *χ*^2^(1) = 35.15, *p <* .001), A-A’ (*β* = 0.76, *χ*^2^(1) = 53.12, *p <* .001), and A-B’ (*β* = 0.51, *χ*^2^(1) = 33.40, *p <* .001). Thus, these results support the prediction that performance would be enhanced with increased encoding for all three of the standard forced-choice test formats. Moreover, we found that the effect of the test-format-contrast was significant within both repetition conditions (see Supplemental Material). In conclusion, we found a significant effect of repetitions, a A-X *>* A-A’ *>* A-B’ test format effect, and a significant interaction. These results are consistent with previous studies and support Predictions 1-3 from MINERVA 2.

Next, we determined if the results from the behavioral experiment supported the predictions of MINERVA 2 within our experimentally-manipulated encoding variability conditions. Here, we compared performance on A-B’ trials for which there was an unequal number of repetitions for the A and B items during learning (i.e., A3x-B’1x, A1x-B’3x) to those for which both the A and B items were presented with 1-repetition (A1x-B’1x) and 3-repetitions (A3x-B’3x). Previous work has been mixed with respect to whether repetitions would enhance (Loiotile & Courtney, 2015; Zhang & Hupbach, 2020) or impair (e.g., Reagh and Yassa, 2014) memory performance. In contrast, results from eye-tracking (Rollins et al., 2019) and EEG (Rollins et al., 2024) have been used to support predictions from computational models that encoding variability may help explain differences in performance between test formats. Thus, the key predictions from MINERVA 2 were that performance would be better when targets (A) were presented with 3-repetitions than 1-repetition (Predictions 1-6) and when the original version of the non-corresponding lure (B) was presented with 1-repetition than 3-repetitions (Predictions 6-8), all of which were borne out in our results (note: all of the *β*’s are positive, indicating that the results follow the expected pattern, and all *ps <* .05; see MINERVA 2 predictions and results above). Here, to compare between the encoding variability conditions (i.e., Predictions 4-8), we defined our models as follows:

~~~
model <-mixed(ProportionCorrect ∼ ExpectedPerf + (1 | ParticipantID),
   data=data, method=‘LRT’, family=binomial, weight=data$n)
~~~

where `ExpectedPerf` is a contrast (+0.5 for the condition with better performance and -0.5 for the condition with worse performance), the model includes random intercepts for participants (more complex random effects models failed to converge), and all other variables are the same as above.

Finally, we tested the prediction from global matching models that target-lure similarity would influence performance across all of the test formats (e.g., related to previous modeling and empirical work with the A-B’ test format: Huffman & Guan, 2025), and we found that the similarity contrast was significant in all of the test formats with similar lures (A1x-A’, A3x-A’, A1x-B’, A3x-B’, A3x-B’1x, A1x-B’3x; all *ps <* .0001; see Supplemental Material and Supplemental Figure 2).

Taken together, the results of the behavioral experiment support the predictions of MINERVA 2 and build on our understanding of the role of encoding variability on human memory. Moreover, our results help resolve a longstanding debate in the field regarding the possible enhancing vs. interfering role of repetitions on lure discrimination. When the target and lure pairs are matched for repetitions, more repetitions enhance memory. In contrast, in conditions in which the original lure item was presented multiple times and the target once, the model and humans are more likely to select that item than in other conditions (e.g., more likely to select B’ in A1x-B’3x than A1x-B’1x, A3x-B’3x, and A3x-B’1x), which suggests that increased matches to the memory trace for the B’, driven by encoding variability, cause higher false alarms (for related results within a signal-detection analysis, see: Zhang & Hupbach, 2020).

Thus, these results provide converging evidence with eye-tracking (Rollins et al., 2019; Rollins, Parks, et al., 2023), EEG (Rollins et al., 2024), and computational modeling (the results here; also, Huffman & Guan, 2025; Huffman & Stark, 2017) results to further our understanding of the role of encoding variability on human memory performance, including better understanding the nature of performance on the A-B’ test format. Our results are also strongly at odds with the competing theory of Competitive Trace Theory (Reagh & Yassa, 2014; Yassa & Reagh, 2013), thus adding to other studies that have suggested that lure discrimination is improved with repetition (Loiotile & Courtney, 2015; Zhang & Hupbach, 2020). Our results are also not consistent with recall-to-reject accounts of memory, which propose that similar lures are rejected following recollection of contextual details associated with the stored memory representation (Rotello & Heit, 1999, 2000). If participants relied on recall-to-reject processing, they would have been expected to demonstrate better performance in A1x-B’3x than A1x-B’1x; however, we observed the opposite pattern of results. We acknowledge that although our results support all of MINERVA 2’s predictions and thus suggest that encoding variability, in terms of the number of features that are encoded for each stimulus, influences mnemonic discrimination, there are potentially other mechanisms that could generate similar predictions (e.g., memory strength). Altogether, our results further support the global matching models framework for the MST (Huffman & Guan, 2025; Huffman & Stark, 2017; Rollins, Huffman, et al., 2023), including accounting for the effects of repetition, experimentally-induced encoding variability, test format, and target-lure similarity (Figure 2 and Supplemental Figure 2).

## Acknowledgements

We thank the Psychology Department at Colby College and Christopher Newport University for helping support this work. We would also like to thank Emily Smith for her assistance with study design and data collection.

## LLM disclosure

We did not use large-language models (LLM’s) or any other generative AI software for any purpose in this work (i.e., we wrote everything ourselves).

## Supplemental Material

### Additional details about sample size determination and our participant pool

We determined our sample size using a power analysis based on the findings from previous research with the A-A’ test format comparing 3 vs. 1 repetitions (Loiotile & Courtney, 2015). Here, we used the function `pwr.t.test` from the pwr package in R with power=0.9 and a significance level of .001, which revealed that we needed to have a minimum sample size of 10 participants. Given that we were additionally interested in testing several novel test formats here, we aimed to collect data from at least 50 participants, thus our sample of 66 participants is greater than our minimum a priori threshold.

The university’s Institutional Review Board approved all procedures before data collection began. Participants were recruited from the Psychology Department’s participant pool and received course credit for participation. We excluded 11 additional participants from the analysis due to technical errors (*n*=7), change in instructions (*n*=3), or failure to meet the inclusion criteria (*n*=1).

### Predictions from Competitive Trace Theory

Here, we also compared the predictions of MINERVA 2 with the predictions from Competitive Trace Theory (CTT). Consistent with Prediction 1 from MINERVA 2, CTT would predict enhanced recognition of a target with multiple exposures when paired with a foil (i.e., better performance on A3x-X vs. A1x-X; Reagh & Yassa, 2014). CTT would also predict better performance when the original version of the non-corresponding lure (B) was presented with 1-repetition than 3-repetitions (Predictions 7-8; i.e., better performance on A1x-B’1x vs. A1xB’3x and A3x-B’1x vs. A3x-B’3x). However, contrary to MINERVA 2, CTT would predict poorer performance on A3x-A’ vs. A1x-A’ and A3x-B’3x vs. A1x-B’1x (Predictions 2-3) because 3-repetitions would lead to semantization, thus resulting in a loss of episodic detail and a higher likelihood of selecting the non-corresponding lure (Reagh & Yassa, 2014; Yassa & Reagh, 2013). Because CTT is a verbal theory (Reagh & Yassa, 2014; Yassa & Reagh, 2013), it was difficult to know exactly what the model would predict for the MINERVA 2 Predictions 4-6, thus again highlighting the importance of computational modeling for “open theorizing” and allowing the generation of testable predictions based on a given set of assumptions (Guest & Martin, 2021).

### Methodological details for the simulations with MINERVA 2

We have previously shown that MINERVA 2 can account for a variety of findings with the Mnemonic Similarity Task, including the test-format effect (Huffman & Guan, 2025; Huffman & Stark, 2017), the effect of target-lure similarity (Huffman & Guan, 2025), and age-related differences observed during childhood (Rollins, Huffman, et al., 2023) and healthy aging (Huffman & Stark, 2017). Thus, we generated predictions of performance across the following test formats: A1x-X, A3x-X, A1x-A’, A3x-A’, A1x-B’1x, A3x-B’3x, A3x-B’1x, and A1xB’3x. Briefly, MINERVA 2 assumes that memories are stored as distributed representations of features, here, modeled as -1, 0, and +1 (in our case, we simulated equal probability of each of these 3 features for each element of the item vector and we used a vector length of 20 features, as in Huffman & Guan, 2025; Huffman & Stark, 2017). MINERVA 2 further assumes that the features are encoded with a given probability representing the learning rate, *L* (here, each feature was encoded with probability of *L* and not encoded [i.e., set to 0] with probability 1 - *L*), which we set to 0.7. We note that in our previous work, we used a learning rate of 0.65 (Huffman & Guan, 2025; Huffman & Stark, 2017), but we increased *L* slightly here to accommodate both the longer list length and the variable list strength here. We generated similar lures as in past work by setting the probability of changing a feature parameter, δ, to 0.16 (Huffman & Guan, 2025; Huffman & Stark, 2017). MINERVA 2 is a multiple-trace model, which assumes that each experience adds a new trace or row to a memory vector (i.e., the columns represent the features and the rows represent the events). For our empirical study, we included 25 stimuli within each condition. Thus, here, we simulated 25 stimuli with one repetition and 25 stimuli with 3 repetitions. On the back end, we then generated and arranged the test items to allow for all of our stated test formats (see above). We ran the model on 1,000 simulated participants.

For the testing phase, we used the standard equations from the original MINERVA 2 model (Hintzman, 1984, 1988). First, for each probe item in a simulated trial, we calculated the similarity of the probe to the memory matrix, *s*_*i*_, using the following equation (Hintzman, 1984, 1988):

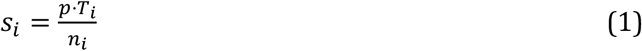

where *p* is the probe item, *T*_*i*_ is one of the items of the memory matrix (i.e., row *i* of the memory matrix, *T*), and *n*_*i*_ is the number of nonzero features in either *p* or *T*_*i*_ (i.e., the similarity function is a normalized dot product). We then calculated the activation, *a*_*i*_, for each of the items with the following equation (Hintzman, 1984, 1988):

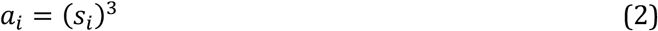

We then calculated the global match, *g*, or summed similarity of the probe item to the entire memory matrix using the following equation (Hintzman, 1984, 1988):

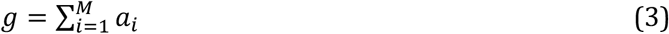

where *M* indicates the number of items in the memory matrix (i.e., then number of rows). For each trial, we calculated the global memory match, *g*, for each of the two simulated probe items. We then calculated the performance using the following equation (cf. Hintzman, 1988; Huffman & Guan, 2025; Huffman & Stark, 2017; Rollins, Huffman, et al., 2023):

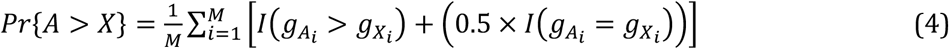

where *M* is the number of items in the memory matrix, *I*() is the indicator function that outputs 1 if the condition is true and zero otherwise, *g*_*Ai*_ is the global match in response to the target item on trial *i*, and *g*_*Xi*_ is the global match in response to the distractor item (either an unrelated foil or a similar lure item) for trial *i*. Note that the left side of the equation calculates the sum of trials in which the global match, *g*, is greater for the target item than the lure item and the right side of the equation simulates random guessing for trials in which the global match is the same for the target item and the lure item (i.e., on average, 50% of these would be correct, hence the 0.5 term here). For our follow up analysis in which we analyzed the data within our binomial mixed model framework, we instead randomly selected the correct vs. incorrect item on a trial-by-trial basis (because the binomial model requires an exact possible proportion of correct trials for each condition).

### Follow-up analyses on the test format effect in each repetition condition

As we reported in the main manuscript, we observed a significant effect of the repetitions contrast (*β* = 0.74, *χ*^2^(1) = 113.70, *p <* .001), the test-format contrast (*β* = 0.91, *χ*^2^(1) = 102.15, *p <* .001), and a repetitions-contrast × test-format-contrast interaction (*β* = 0.20, *χ*^2^(1) = 5.67, *p* = .017). To further explore the test format effects, we conducted follow-up analyses to test whether the test-format effect was significant within each repetition condition. We defined our models as follows:

~~~
model <-mixed(ProportionCorrect ∼ TestFormat + (1 | ParticipantID),
   data=data, method=‘LRT’, family=binomial, weight=data$n)
~~~

where `ProportionCorrect` is the proportion correct for a given format, `TestFormat` is a contrast of expected performance based on MINERVA 2 (+1 for A-X, 0 for A-A’, -1 for A-B’), `(1 | ParticipantID)` is the random effect of participant (here we used random intercepts because more complex models failed to converge), and `weight=data`$`n` is a vector of the number of trials within each condition (here, we had 25 trials for each of these conditions). We found a significant effect of the test-format contrast within the 1-repetition data (*β* = .71, *χ*^2^(1) = 204.46, *p <* .001). Likewise, the planned comparisons revealed that all 3 test formats were significantly different than each other within the 1-repetition data: A1x-X *>* A1x-A’ (*β* = 1.10, *χ*^2^(1) = 98.79, *p <* .001), A1x-X *>* A1x-B1x’ (*β* = 1.54, *χ*^2^(1) = 216.75, *p <* .001), and A1x-A’ *>* A1x-B1x’ (*β* = 0.43, *χ*^2^(1) = 24.20, *p <* .001). Altogether, these results support the predictions of MINERVA 2 that there will be a A-X *>* A-A’ *>* A-B’ test format effect within the 1-repetition condition (compare these results with Figure 2A). Next, we tested whether the test format-effect was also significant within the 3-repetition condition by running a similar analysis on the data from the 3-repetition condition. We found a significant effect of the test-format contrast for the 3-repetition condition (*β* = 0.91, *χ*^2^(1) = 203.23, *p <* .001). Likewise, the planned comparisons revealed that all 3 test formats were significantly different than each other within the 3-repetition data: A3x-X *>* A3x-A’ (*β* = 1.12, *χ*^2^(1) = 63.87, *p <* .001), A3x-X *>* A3x-B3x’ (*β* = 1.98, *χ*^2^(1) = 211.65, *p <* .001), and A3x-A’ *>* A3x-B’3x (*β* = 0.69, *χ*^2^(1) = 42.16, *p <* .001). Altogether, these results support the predictions of MINERVA 2 that there will be a A-X *>* A-A’ *>* A-B’ test format effect within the 3-repetition condition (compare these results with Figure 2A). In summary, we found evidence to support the A-X > A-A’ > A-B’ test format effect in both the 1-repetition and 3-repetition condition, thus building on previous behavioral and modeling findings with the typical 1-repetition version of the MST (Huffman & Guan, 2025; Huffman & Stark, 2017; Rollins et al., 2019, 2024; but see Migo et al., 2009).

### Analysis of the data from the MINERVA 2 simulations

Following the somewhat unexpected finding of the interaction in the empirical data (see main text), we went back to our original a priori modeling results and decided to analyze the data from the model’s predictions (i.e., we had previously only visually interpreted the data to note the prediction of the test format effect and the repetition effect) using the original modeling parameters. We used the same mixed-effects modeling approach for the simulated data (except we only included random intercepts for participants here, since more complex random effects models failed to converge). Here, we found that MINERVA 2 made qualitatively similar predictions to the empirical results: we observed a significant effect of the repetitions contrast (*β* = 1.44, *χ*^2^(1) = 6,044.28, *p <* .0001) and the test-format contrast (*β* = 1.04, *χ*^2^(1) = 9,354.52, *p <* .0001). Moreover, we observed a significant repetitions-contrast × test-format interaction (*β* = .80, *χ*^2^(1) = 1,148.13, *p <* .0001). Furthermore, consistent with the empirical data, follow-up pairwise comparisons of estimated marginal means (using emmeans within R; here we used a categorical version of the Repetitions and Test Format for easier interpretation of the effects) revealed that the repetition effect was (numerically) largest for the A-X format (odds ratio = 59.88, *p* < .0001) followed by the A-A’ format (odds ratio = 2.96, *p* < .0001) and smallest for the A-B’ format (odds ratio = 59.88, *p* < .0001). Likewise, as in the empirical data, these findings are further supported by the positive *β* for the interaction term, thus suggesting the prediction of an interacting benefit of increased repetitions with the increasing performance across the test format contrast. We note that the overall magnitude of effects are larger for the simulations, but we included 1,000 simulated participants and we would like to emphasize that we were concerned with qualitative effects (i.e., rather than attempting to quantitatively fit the data).

### Lure Bins

We next tested whether performance varied as a function of target-lure similarity across all of the test formats with similar lures. Here, we used the “lure bins” based on behavioral performance from Stark et al. (2013). Specifically, Stark et al. (2013) created the “lure bins” from the most similar in lure bin 1 (highest “old” responses) to the most dissimilar in lure bin 5 (the lowest “old” responses). Here, we set up a contrast for lure bins 1 to 5: [-2, -1, 0, 1, 2] (the `LureBinContrast` variable in the equation below), which allowed us to test our specific prediction that performance would be lowest for lure bin 1 and progressively better through lure bin 5. We again ran our analysis within afex with a binomial family in R, using the following model:

~~~
model <-mixed(ProportionCorrect ∼ LureBinContrast + (1 | ParticipantID),
   data=data, method=‘LRT’, family=binomial, weight=data$n)
~~~

Here, we had 5 trials within each lure bin for each test type. We ran this model separately within each condition and repetition to determine whether there was a significant effect of lure bin, which would be consistent with MINERVA 2 (Huffman & Guan, 2025). We found a significant effect of the lure bin contrast in all of the conditions: A-A’ 1x (*β* = 0.39, *χ*^2^(1) = 68.19, *p <* .001), A-A’ 3x (*β* = 0.64, *χ*^2^(1) = 94.14, *p <* .001), A-B’ 1x (*β* = 0.29, *χ*^2^(1) = 47.44, *p <* .001), A-B’ 3x (*β* = 0.38, *χ*^2^(1) = 58.99, *p <* .001), A3x-B’1x (*β* = 0.62, *χ*^2^(1) = 63.51, *p <* .001), and A1x-B’3x (*β* = 0.44, *χ*^2^(1) = 107.97, *p <* .001; note: the original model failed to converge, so we dropped the lure bin 3 condition, in which case the model converged, thus our model was slightly different here, but included the contrast: [-1.5, 0.5, 0.5, 1.5]). Thus, the effect of lure bin is consistent across all 6 test formats with similar lures. Moreover, our results support recent work with several global matching models, including MINERVA 2, which all predicted that performance on test formats with targets and similar lures would vary as a function of the target-lure similarity (Huffman & Guan, 2025).

---

**Supplemental Figure 1:**
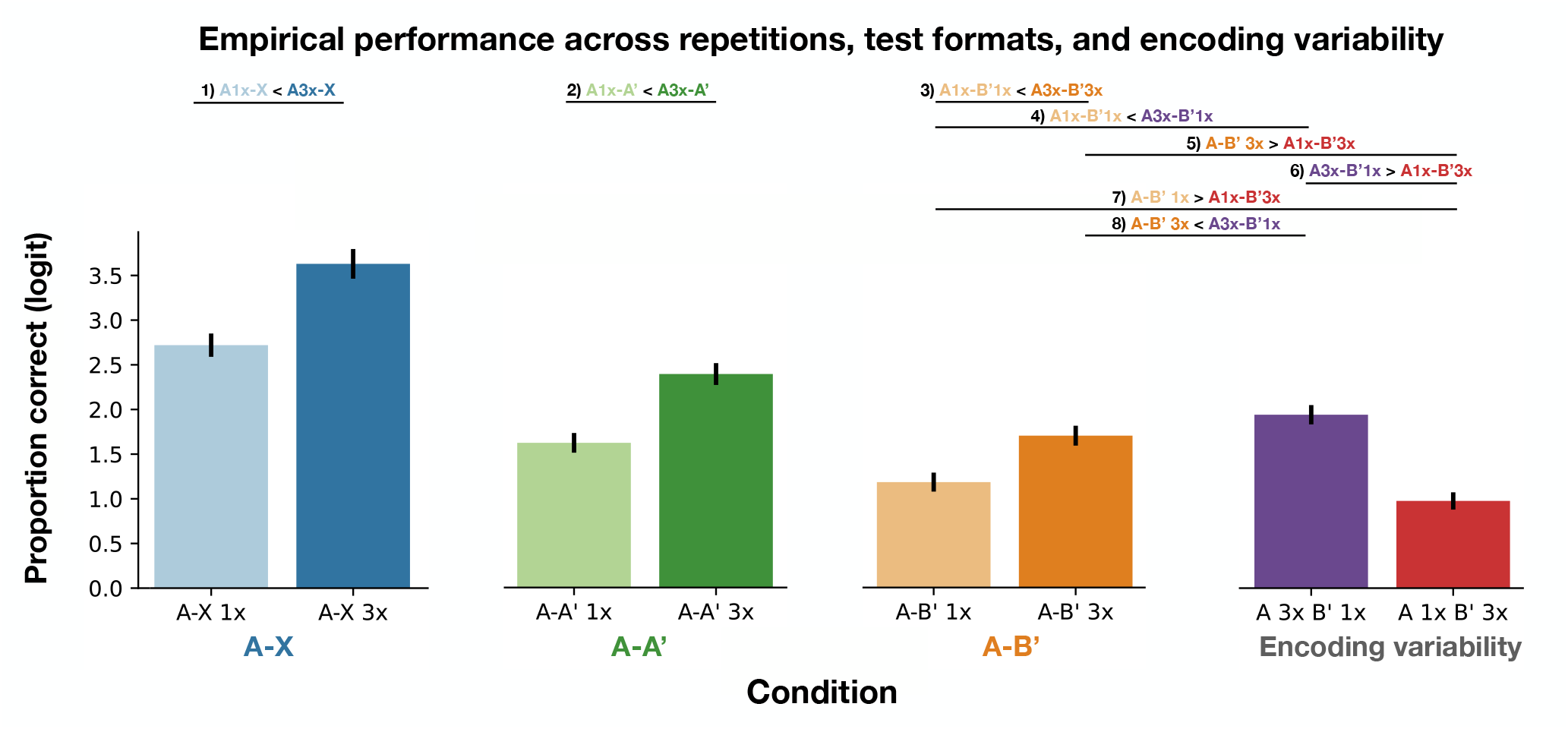
Here, we replotted the data from Figure 1B on the logit scale (i.e., the original scale as our statistical analysis with the mixed-effects binomial models). The repetition-contrast × test-format-contrast format interaction effect for the A-X, A-A’, and A-B’ test format is clearer on this scale: the repetition effect was (numerically) largest for the A-X test format, followed by the A-A’ test format, and smallest for the A-B’ test format. The data depict the estimated marginal means and the 95% standard errors from our binomial models.

**Supplemental Figure 2:**
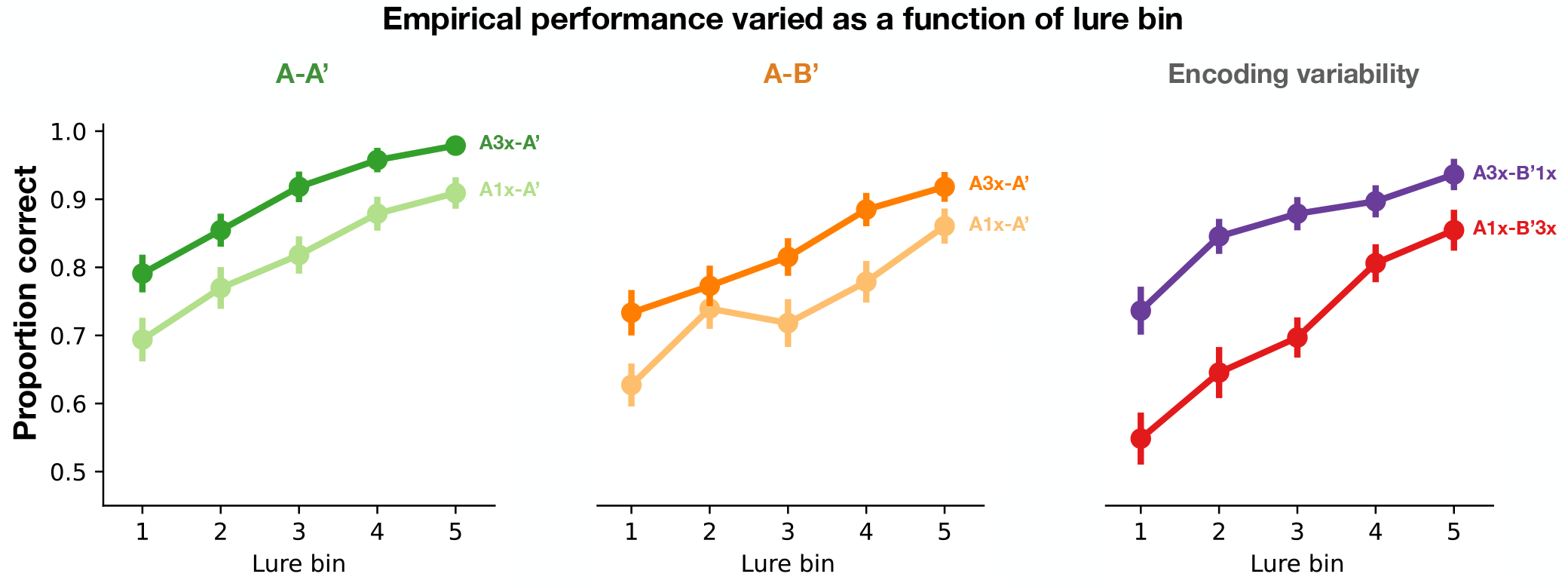
Performance varied as a function of the lure bin across all 6 of the force-choice conditions with similar lures (i.e., a significant positive effect of the lure-bin contrast of [-2, -1, 0, +1, +2] across lure bins 1-5, which are ordered from most similar to least similar, based on previous research [see above]; all *ps <* .001). The data for the plots indicate the raw means and 95% standard errors but we used a binomial model for analysis.

